# A novel Versatile Peroxidase from Lentinus squarrosulus (MH172167)towards enhanced delignification and digestibility of crop residues

**DOI:** 10.1101/394098

**Authors:** R Aarthi, Ramya G Rao, Vandana Thammaiah, SM Gopinath, Manpal Sridhar

## Abstract

Scarcity of quality feed is a major constraint concerning livestock productivity with recalcitrant lignin hindering utilization of crop residues as quality animal feed. Degradation of lignin in nature is contributed by white-rot fungi through their enriched ligninolytic system. Versatile Peroxidase plays a key role in ligninolysis through its capability to oxidize diverse class of aromatics without mediators. In this study, wild isolates of wood rotting fungi were screened for potential peroxidases oxidizing manganese and aromatic compounds. The strain identified as *Lentinus squarrosulus* (TAMI004, BankIt2098576 MH172167) was monitored for enzyme activity in solid state and submerged fermentation. *L. squarrosulus* demonstrated predominant Versatile Peroxidase activity amongst the screened wild isolates displaying hybrid characteristic of manganese oxidation and manganese independent reactions on aromatic compounds. The manganese oxidizing peroxidase activity evidenced in submerged fermentation was 12 IU/L whereas in solid state fermentation it was 131 IU/L. This ability to act through manganese mediated and independent reactions on phenolics reveals its biotechnological and industrial significance. Treatment of common crop residues with crude extract of *L. squarrosulus* rich in Versatile Peroxidase obtained from both Solid state and submerged fermentations showed a decrease in their Neutral Detergent Fiber, Acid Detergent Fiber and Acid Detergent Lignin content showing biodegradation, substantiating the ligninolytic ability and more prominently increase in their digestibility. To the best of our knowledge, this is the first report describing Versatile Peroxidase from *Lentinus squarrosulus* with potential to augment the ruminant digestibility of crop residues.

**Importance:** Versatile Peroxidase of White-rot fungi, a relatively less studied lignolytic enzyme, is very efficient in depolymerization of lignin macromolecule through its multivalent catalytic sites. Lignin degradation is very appealing from the application perspective as attack on lignin exposes the energy affluent polysaccharides for utilization in extensive biotechnological applications. Reports on relevance of Versatile Peroxidase for these purposes are still emerging, however the role of ligninolytic enzymes especially Versatile Peroxidase in enriching ruminant feed is yet unturned. Here, this work demonstrates the potential of Versatile Peroxidase from a novel species *Lentinus squarrosulus* in delignification thereby upgrading the digestibility and nutritive value of crop residues. The observations validate the importance of the enzyme in improvement of crop residues for feeding ruminants in the current scenario where, livestock productivity is severely impacted by lack of quality feed and demand for alternate feed resources is intensifying.

## Introduction

World in this phase of phenomenal growth requires economical and effective technological applications in every aspect to reinforce its development. Sustainable energy and waste management are the pillars of socio-economic development. Likewise, environmental health is equally important besides promoting socio-economic prosperity. On these lines, countries worldwide are witnessing an increase in demand for food and fuel. This urges us to look for alternate sources of feedstock for fuel without compromising food security. Sustainable agricultural systems are seen as drivers of food security and environmental viability. Here, livestock plays an inexorable role in supporting sustainable agriculture. However, the quest for quality feed for the expanding livestock community is still a concern impacting the efficiency of livestock production. One of the keys to disengage these challenges lies in the lignocellulosic biomass generated through cropping practices and as by-products of agro-industries worldwide (1). Interestingly, utilization of lignocellulosic biomass also addresses the issue of residue disposal which in current times is a major issue contributing to greenhouse gases emission. These lignocellulosic residues are pertinent in diverse applications of biofuel production, paper pulp manufacture, animal feed to mention a few. Having said that, exploitation of these lignocelluloses relies heavily on cost effective destruction of the biomass. Biomass deconstruction is necessitated owing to the recalcitrance of the lignocellulosic complex emerging from crystallinity of cellulose embedded in the lignin-hemicellulose complex, rigidity and hydrophobicity of lignin. Although, diverse categories of pretreatment methods have their own advantages, biological method of degradation stands out economically. A fascinating group of fungi termed “White-rot” dominates the biological lignin degraders by its competence and rich network of ligninolytic enzymes. White-rot fungi surpass its affiliates from bacteria and other fungi of different ecological group through these unique lignolytic capabilities of selective lignin degradation (2). Not surprisingly, there is a spurt in research in recent past on the characterization of these ligninolytic enzymes for their ability to disintegrate lignocellulosic complex in a way amenable to downstream biotechnological applications. Ligninolytic system of these white-rot fungi comprises mainly of Laccase, Manganese Peroxidase, Lignin Peroxidase and Versatile Peroxidase besides other enzymes involved in peroxide generation (3). Laccase, the phenol oxidase and the heme peroxidases viz. Manganese Peroxidase, Lignin Peroxidase oxidize the non-phenolic units of lignin through mediators while the phenolic units are attacked directly. Extensive research has established the role of these enzymes in lignin degradation. Versatile Peroxidase or VP (EC 1.11.1.16) is unique in the system for its ability to oxidize the phenolic and non-phenolic compounds free of redox mediators. This is feasible owing to the hybrid architecture of Lignin Peroxidase and Manganese Peroxidase being present in Versatile Peroxidase and adaptation of amino acid residues in the catalytic environment favorable for high potential substrate oxidation (4). For this reason, Versatile Peroxidase oxidizes high molecular weight substrates through catalytic tryptophan at the surface resembling Lignin Peroxidase and Mn^2+^ to Mn^3+^ as in Manganese Peroxidase (5). Nevertheless, there is dearth of studies on Versatile Peroxidase as this enzyme exhibits considerable complexity for identification among the other ligninolytic peroxidases. This study focuses on screening of a novel Versatile Peroxidase from wild isolates with exceptional efficiency for aromatics oxidation besides oxidation of manganese.

## Materials and Methods

### Chemicals

Chemicals used in the study were of analytical grade unless otherwise stated. Reactive Black 5 (RB5), 2, 2’-azino-bis (3-ethyl benzothiazoline-6-sulfonic acid) (ABTS) and 2, 6 dimethoxyphenol (DMP) used for assay were procured from Sigma Aldrich (USA).

### Culture Propagation and Screening

Twenty isolates of wood rotting fungi from humid, tropical regions of Western Ghats (12.9763° N, 77.5929° E, 8.9339° N, 77.2780° E) were brought to pure cultures by aseptic propagation of inner tissue of pileus/stipe of basidiocarp on potato dextrose agar slants at 28ºC (6). All the isolates were appraised for their ligninolytic ability through Bavendamm test of oxidation of 30mM gallic acid and 3mM tannic acid in discrete reactions (7). Additionally, the enzymatic reactivity of the isolates towards guaiacol, the O-methoxyphenol at a concentration of 8mM on solid media confirmed their phenolic oxidation ability. The strains with positive reactions to the above analysis were selected for the screening of Versatile Peroxidase production.

Subsequently, agar plates supplemented with 100µM Reactive Black 5, an azo dye were observed for oxidative decolouration. The plates were inoculated with 4mm mycelial plug from freshly grown culture and observed at regular intervals for halo of decolorization. Reactive Black 5 is a heavily recalcitrant double azo dye possessing high redox potential and hence chosen for the present study. Decolorization of sulfonphthaleine dyes Bromophenol blue, Bromothymol blue, Bromocresol green were studied by supplementing liquid culture medium with 0.01% of the corresponding dye (8).

### Culture Conditions

The basal medium for cultivation comprised per liter Glucose 10 g, NH_4_H_2_PO_4_ 2 g Yeast extract 2 g and trace element solution 1 ml. Trace elements include MgSO_4_ 3 g/L, CuSO_4_ 0.005 g/L, ZnSO_4_ 0.1 g/L, FeSO_4_ 0.1 g/L, CaCl_2_ 0.05 g/L. Fungal cultures pre-grown in glucose-ammonium medium were used as inocula. Prior to inoculation, the cells were homogenized and added to the production medium. Flasks were maintained under continuous agitation of 120 rpm. Solid state fermentation was implemented using pre-treated poplar wood chips moistened with basal medium (9). 250 ml Erlenmeyer flask containing 5 g of pre-treated wood chips layered with 25 ml basal medium was inoculated with homogenized mycelia from freshly grown culture. Cultures were incubated at 28ºC in dark. Enzyme analysis was performed using extracellular fluids from submerged and solid-state fermentation. Effect of supplementation of different concentrations of manganese (0 – 500µM) on enzyme production was analyzed by addition of appropriate quantities of manganese as MnSO_4_ to the basal medium. All experiments were performed in triplicates.

### Decolorization studies

To ascertain the rate of degradation of the azo dye Reactive Black 5, liquid medium supplemented with 100 µM dye was inoculated with 4 mm mycelial plug from freshly grown culture and monitored for decolorization spectrophotometrically at the absorbance maxima of the dye once in 24 hours. Flasks were incubated at 28ºC under continuous agitation at 120 rpm. Un-inoculated flask supplemented with dye served as control. Decolorization efficiency was assessed as % decolorization using the below equation

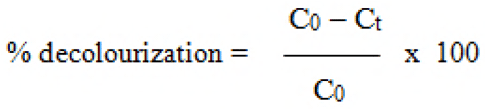

Where *C*_*0*_ is the initial concentration of the dye and *C*_*t*_ is concentration of the dye at *t* hours. All experiments were performed in triplicates.

### Analytical Measurements

Laccase activity was estimated by the oxidation of 1.6 mM ABTS in 100 mM sodium acetate buffer (pH 4.5) at 420 nm (ε_420_ 36000 M^−1^ cm^−1^) (10). Oxidation of ABTS by peroxidases was corrected by subtracting the activity in presence of 0.5µg/ml Catalase. Lignin Peroxidase activity was measured by monitoring the oxidation of 2 mM Veratryl alcohol in 100 mM sodium tartrate buffer pH 3. The formation of veratrylaldehyde was measured at 310 nm (ε_310_=9300M^-1^ cm^-1^). Manganese peroxidase activity was deduced from the formation of Mn^3+^ malonate at 270 nm. The assay mixture consisted 0.5 mM MnSO_4_ and 100 mM malonate buffer pH 4.5 and the reaction was initiated with 0.1 mM H_2_O_2_. Oxidation of RB5 was determined in 100 mM sodium tartrate buffer pH 3 with 10 µM of RB5. The reaction was initiated by addition of 0.1 mM H_2_O_2_ and monitored through decrease in absorbance at 598 nm (ε_598_24000 M^−1^ cm^−1^). Oxidation of 2, 6 dimethoxyphenol (DMP) was determined at 468 nm in a reaction mixture containing 0.1 mM DMP, 100 mM sodium tartrate buffer pH 3 and 0.1 mM H_2_O_2_. One unit of enzyme is defined as the amount of enzyme that transforms 1 µmol of substrate or that forms 1 µmol of product per minute. Absorbance measurements were carried out in Shimadzu UV 1800 spectrophotometer. Reducing sugars were estimated by Dinitrosalycilic acid (DNS) method with D-glucose as standard (11). Microbial biomass was deduced by filtering the media contents through pre-weighed dried Whatman no 1 filter paper followed by overnight drying at 70^°^C.

### Culture Identification

This efficient strain exhibiting ligninolytic activity was identified in a discrete study undertaken by Rao *et al* (12). Genomic DNA from freshly grown fungal culture was isolated by microwave method which then served as a template for PCR with ITS 1 (5′-TCC GTA GGT GAA CCT GCG G-3′) and ITS 4 (5′-TCC TCC GCT TAT TGA TAT G-3′) as primers. The amplified product spanning 18s rDNA, 28s rDNA, 5.8s rDNA along with ITS1 and ITS4 regions was sequenced and the ITS sequence obtained was then subjected to sequence comparison through BLAST nucleotide search tool of NCBI. Low complexity filter was set and the sequences with lowest expect value and maximal identity were selected.

### Proximate Analysis

Five commonly available crop residues such as paddy straw, finger millet straw, foxtail millet straw, little millet straw and barnyard millet straw were milled to 1-2 cm length, dried at a constant temperature of 70±2 ºC for use in biodegradation studies. Crude enzyme produced by the fungus through solid state and submerged fermentation modes was harvested on the peak of enzyme production and exploited for treating the crop residues. The crop residues were treated with the crude enzyme at 40% (v/w) and incubated for 24 hours followed by drying at 70±2 ºC. The samples were then analyzed for proximate principles of NDF, ADF and ADL following the method of Van soest *et al* 1991 (13).

### Digestibility Analysis

Subsequent to proximate analysis, finger millet straw and little millet straw were analyzed for their *in vitro* dry matter digestibility through a two stage *in vitro* method (14). Prior to digestion of the samples, fermentation medium was prepared by adding 2.5 g/L casein hydrolysate and 0.15% resazurine solution to McDougall’s buffer. McDougall’s buffer composed per liter NaHCO_3_ 9.80g, Na_2_HPO_4_ 7g, KCl 0.57g, NaCl 0.47g, MgSO_4_.2H_2_O 0.12g, CaCl_2_ 0.04g. Reducing agent (625 mg cysteine hydrochloride dissolved in 95 ml distilled water, 4ml of 1N NaOH added followed by addition of 625 mg sodium sulphide flakes) was introduced to the fermentation medium and then flushed with CO_2_ to remove the air out of the medium. Rumen contents were obtained from a male cow through a permanent fistula. The contents were maintained at 39°C and carefully filtered through two layers of cheese cloth. 40 ml of fermentation medium with 10 ml of rumen liquor were then added to 0.5 g of finely ground sample. The sample flasks were flushed with CO_2_ and incubated at 39°C for 48 hours with continuous shaking. After incubation, the samples were extracted using neutral detergent solution to assess their digestibility on dry matter basis.

### Statistical Analysis

All treatments were performed in three replicates. Proximate analysis was evaluated using PROC GLM in SAS 9.3 software. Significant differences between treatments were determined using t-test at α value of 0.05.

## Results

### Screening of wild isolates for Ligninolytic enzymes

Oxidation of phenolics was indicated by the formation of brown to reddish brown radiance on agar plates. Strains were considered positive for ligninolytic activity only on oxidation of all the phenolic indicator compounds subjected in the study. Accordingly, 60% of wood rotting strains screened in the present study were positive for ligninolytic activity. Subsequently, 30% of strains were found proficient to decolorize the recalcitrant azo dye Reactive Black 5 (RB5) and were considered apparent for Versatile Peroxidase. Though sulfonphthaleine dyes decolorization was observed in majority of the isolates, the azo dye decolorization was exhibited by markedly few isolates. The typical decolorization paces of the isolates are depicted in Fig 1. The efficient isolate designated as TAMI004 as explored through these preliminary studies was further evaluated in liquid culture for production of ligninolytic enzymes. In addition, microscopic observation of the type of hyphae, septa, clamp connections and the features of the spores confirmed these fungal strains as belonging to basidiomycetous family.

**Fig 1:**
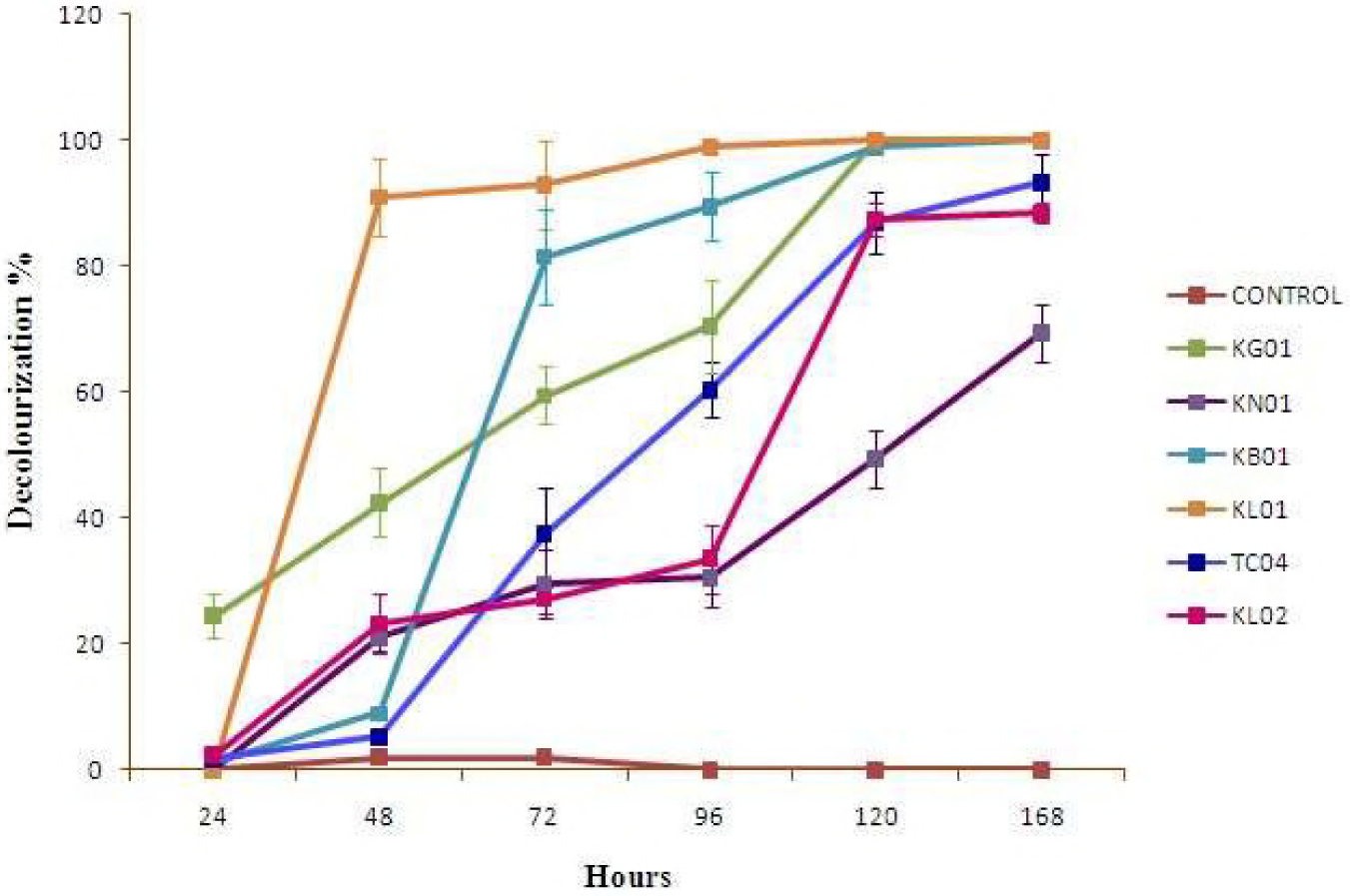
RB5 Decolorization profile of the wild isolates in potato dextrose medium. TC04 refers to *Lentinus squarrosulus*

### Identification of wild isolate

Genomic DNA extracted through microwave method was of highest purity as determined through nanodrop spectrophotometer. Amplification through ITS 1 and ITS 4 primers yielded 700 bp sequence encompassing ITS 1, ITS 2 and rDNA regions and deposited to Genbank with accession nos MH172167, MH172168. Sequence comparison through NCBI BLAST identified the strain as basidiomycete *Lentinus squarrosulus* (TAMI004, BankIt2098576 MH172167).

### Characterization of ligninolytic enzyme production

*Lentinus* species studied here is a rich producer of Laccase as determined by ABTS oxidation ability besides peroxidase activities. The remarkable ability is the decolorization of RB5 and RBBR in manganese independent reactions (Fig 2). Decolorization of sulfonphthaleine dye bromophenol blue is illustrated in Fig 3. In liquid culture, Manganese oxidizing peroxidase activity reached its maximum after 7 days, at the very end of active tropophase (log phase) after which the activity declined. Residual reducing sugar was nearly negligible at this phase (Fig 5). To evaluate the activity of the manganese oxidizing peroxidase enzymes in manganese containing medium, the above isolate was cultivated in basal medium with varying concentrations of manganese from 0 –500µM. Here, the RB5 oxidizing peroxidase activity was predominant in the medium devoid of manganese though manganese oxidizing peroxidase activity was significant in media with 0 and 500 µM manganese. In contrary, only trivial veratryl alcohol oxidation was observed with this isolate. Manganese dependent and independent activities on 2, 6 dimethoxy phenol were also exhibited by this species. The pH optima for RB5 oxidizing activity and manganese oxidizing activity were observed as 3 and 4.5 respectively. Solid state fermentation demonstrated superior ligninolytic ability in comparison to submerged fermentation. The manganese oxidizing peroxidase activity evidenced in submerged fermentation was 12 IU/L whereas in solid state fermentation it was 131 IU/L. The activity values obtained in the current study were comparable to those produced by *Bjerkandera sp* in glucose ammonium medium (15). This *Lentinus* strain also grew profusely in solid state fermentation on wood chips, wood being the preferred substrate of this fungus (Fig 4).

**Fig 2:**
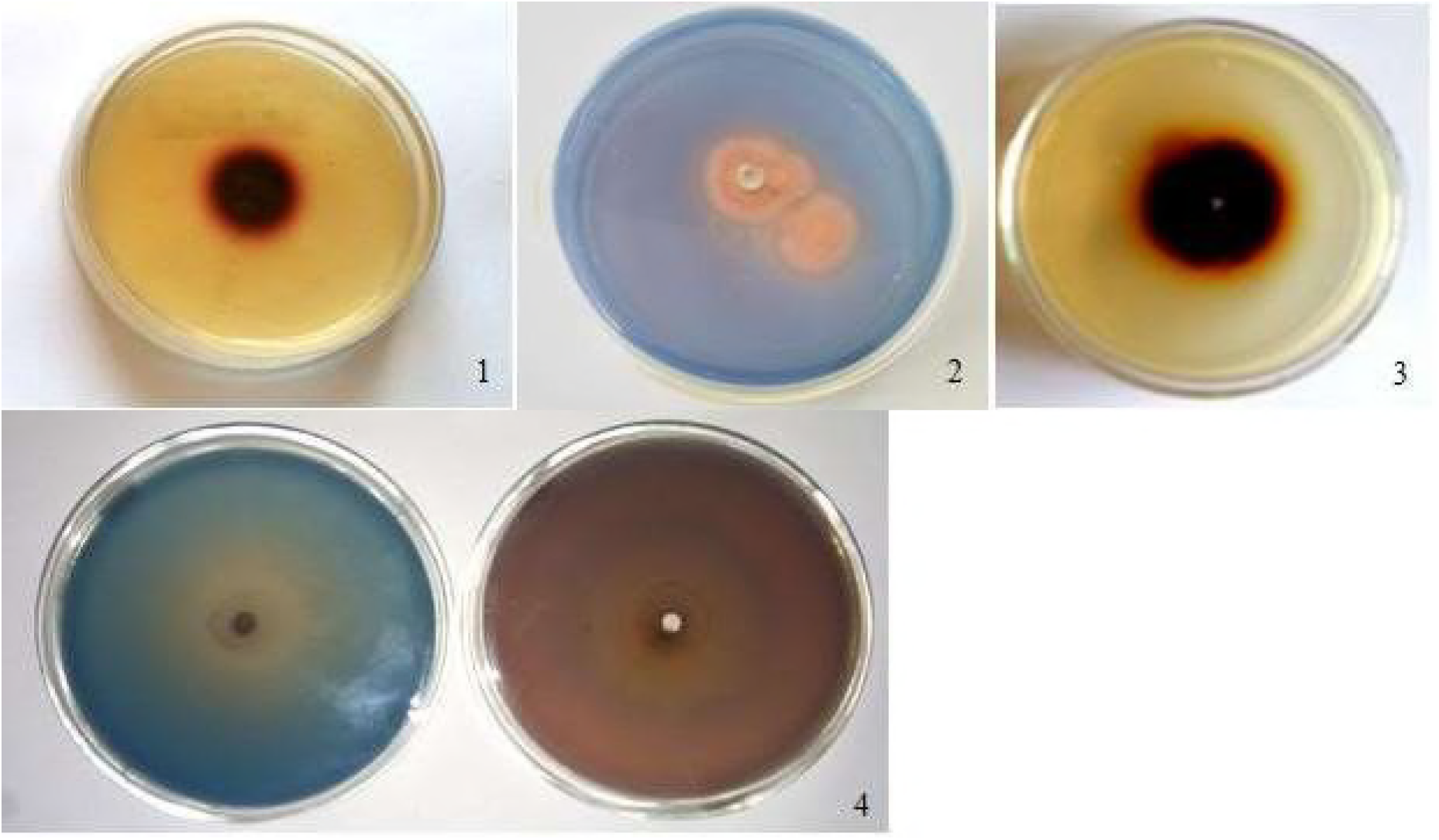
Phenolics oxidation by *Lentinus squarrosulus* (1.) Oxidation of guaiacol (2.) Decolorization of RB5 (3.) Oxidation of gallic acid (4.) Decolorization of RBBR after 3 and 7 days of culture.

**Fig 3:**
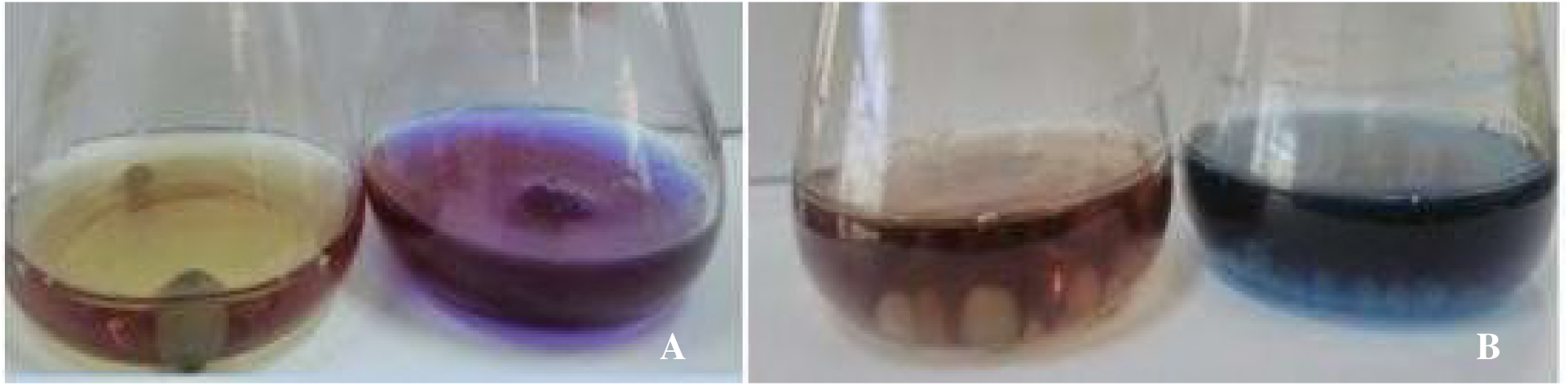
(A) Decolorization of sulfonphthaleine dye Bromophenol blue (B) Decolorization of RB5 in liquid culture in comparison to negative control.

**Fig 4:**
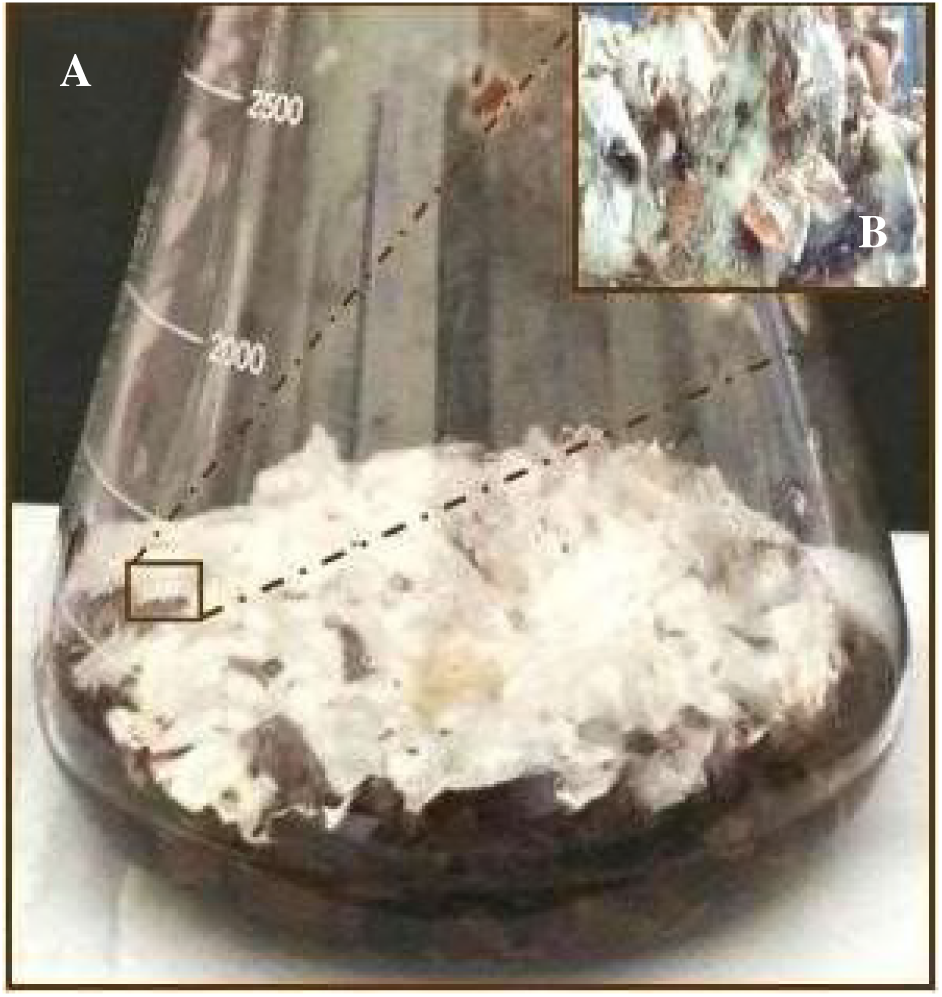
(A) Solid state fermentation of *Lentinus squarrosulus* on wood chips (B) close up view of colonization of the fungus on wood chips.

**Fig 5:**
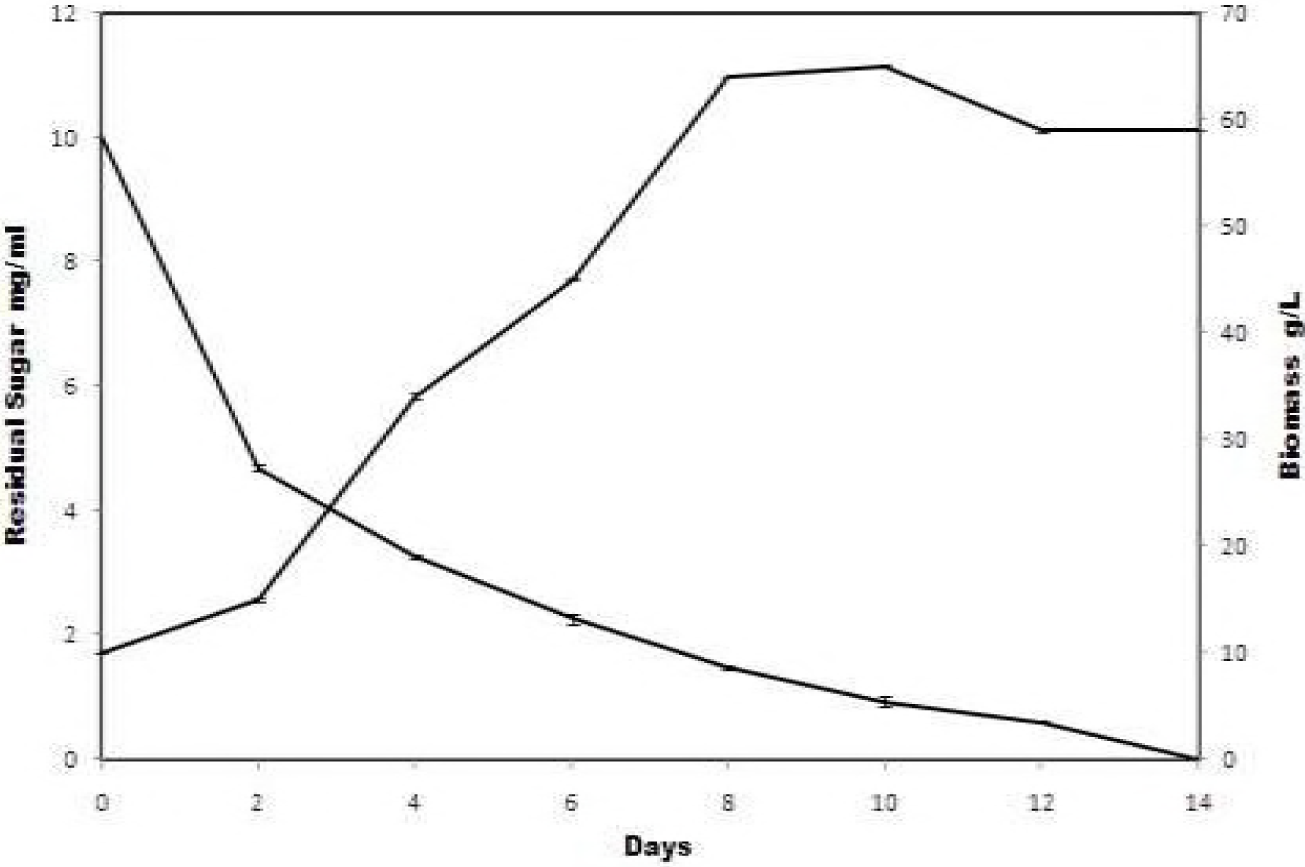
Biomass growth and reducing sugar profile of *Lentinus squarrosulus* during submerged fermentation.

### Estimation of Proximate principles

All straws showed decrease in the Neutral Detergent Fiber (NDF), Acid Detergent Fiber (ADF) and Acid Detergent Lignin (ADL) contents of enzyme treated samples irrespective of the mode of fermentation as evident from the means of the treatments in Table 1 (Fig 7, 8, 9). On the whole, NDF recorded 3.6% and 4.2% reduction in solid state (SSF) and submerged (SmF) fermentation enzyme treatments. There was 5.2% and 2.9% decrease in acid detergent lignin in SSF and SmF enzyme treatments respectively. Little millet, foxtail millet and paddy straw exemplified statistically promising results while finger millet straw and barnyard millet showed slightest response. In addition, it was inferred that solid-state fermentation was comparatively significant to submerged fermentation.

**Table 1:**
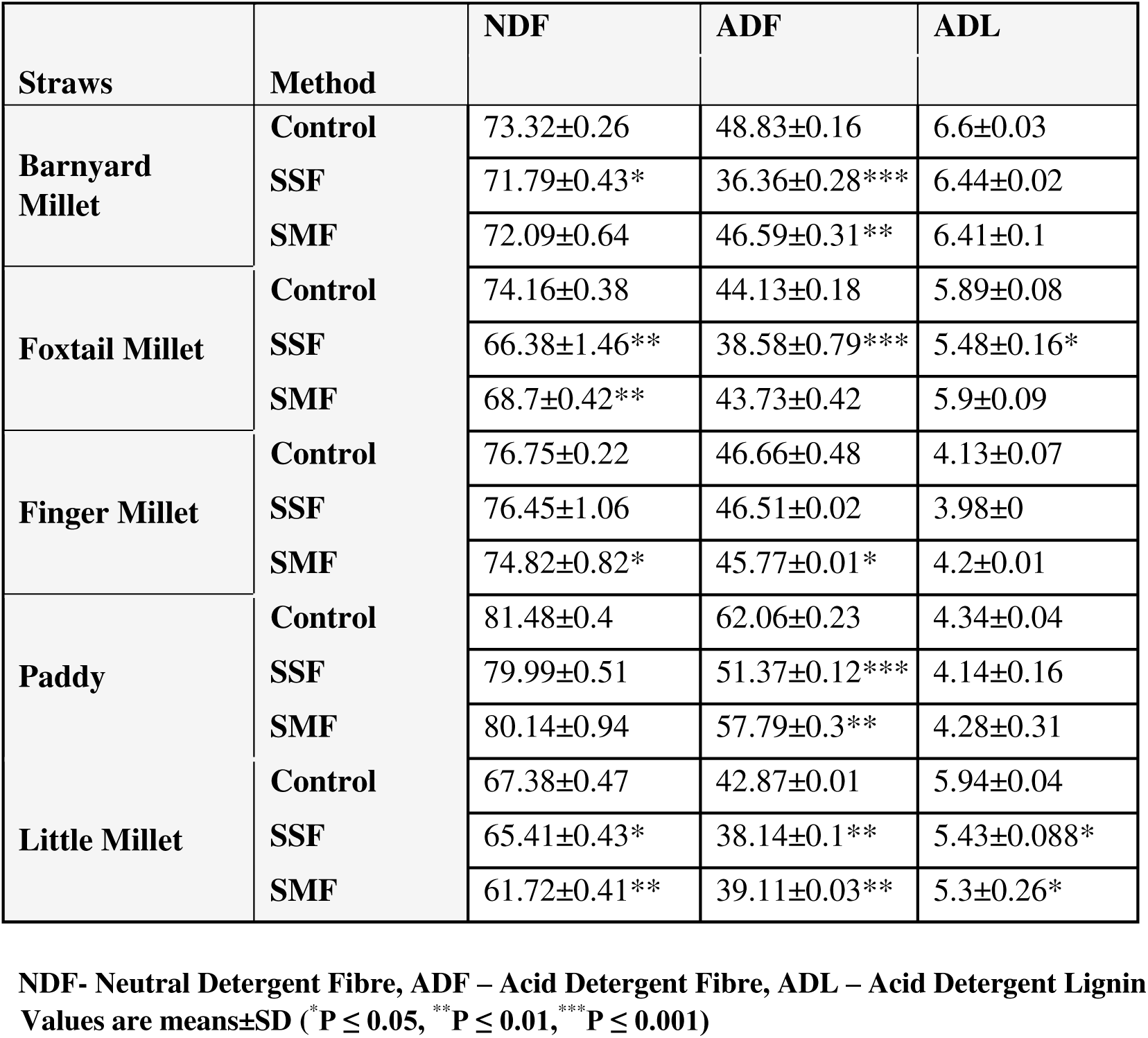
NDF, ADF and ADL of control and enzyme treated straws.

| Straws | Method | NDF | ADF | ADL |
| --- | --- | --- | --- | --- |
| **Barnyard Millet** | **Control** | 73.32±0.26 | 48.83±0.16 | 6.6±0.03 |
|  | **SSF** | 71.79±0.43* | 36.36±0.28*** | 6.44±0.02 |
|  | **SMF** | 72.09±0.64 | 46.59±0.31** | 6.41±0.1 |
| **Foxtail Millet** | **Control** | 74.16±0.38 | 44.13±0.18 | 5.89±0.08 |
|  | **SSF** | 66.38±1.46** | 38.58±0.79*** | 5.48±0.16* |
|  | **SMF** | 68.7±0.42** | 43.73±0.42 | 5.9±0.09 |
| **Finger Millet** | **Control** | 76.75±0.22 | 46.66±0.48 | 4.13±0.07 |
|  | **SSF** | 76.45±1.06 | 46.51±0.02 | 3.98±0 |
|  | **SMF** | 74.82±0.82* | 45.77±0.01* | 4.2±0.01 |
| **Paddy** | **Control** | 81.48±0.4 | 62.06±0.23 | 4.34±0.04 |
|  | **SSF** | 79.99±0.51 | 51.37±0.12*** | 4.14±0.16 |
|  | **SMF** | 80.14±0.94 | 57.79±0.3** | 4.28±0.31 |
| **Little Millet** | **Control** | 67.38±0.47 | 42.87±0.01 | 5.94±0.04 |
|  | **SSF** | 65.41±0.43* | 38.14±0.1** | 5.43±0.088* |
|  | **SMF** | 61.72±0.41** | 39.11±0.03** | 5.3±0.26* |

**NDF- Neutral Detergent Fibre, ADF – Acid Detergent Fibre, ADL – Acid Detergent Lignin** **Values are means±SD (\* P ≤ 0.05, \*\* P ≤ 0.01, \*\*\* P ≤ 0.001)**

**Fig 6:**
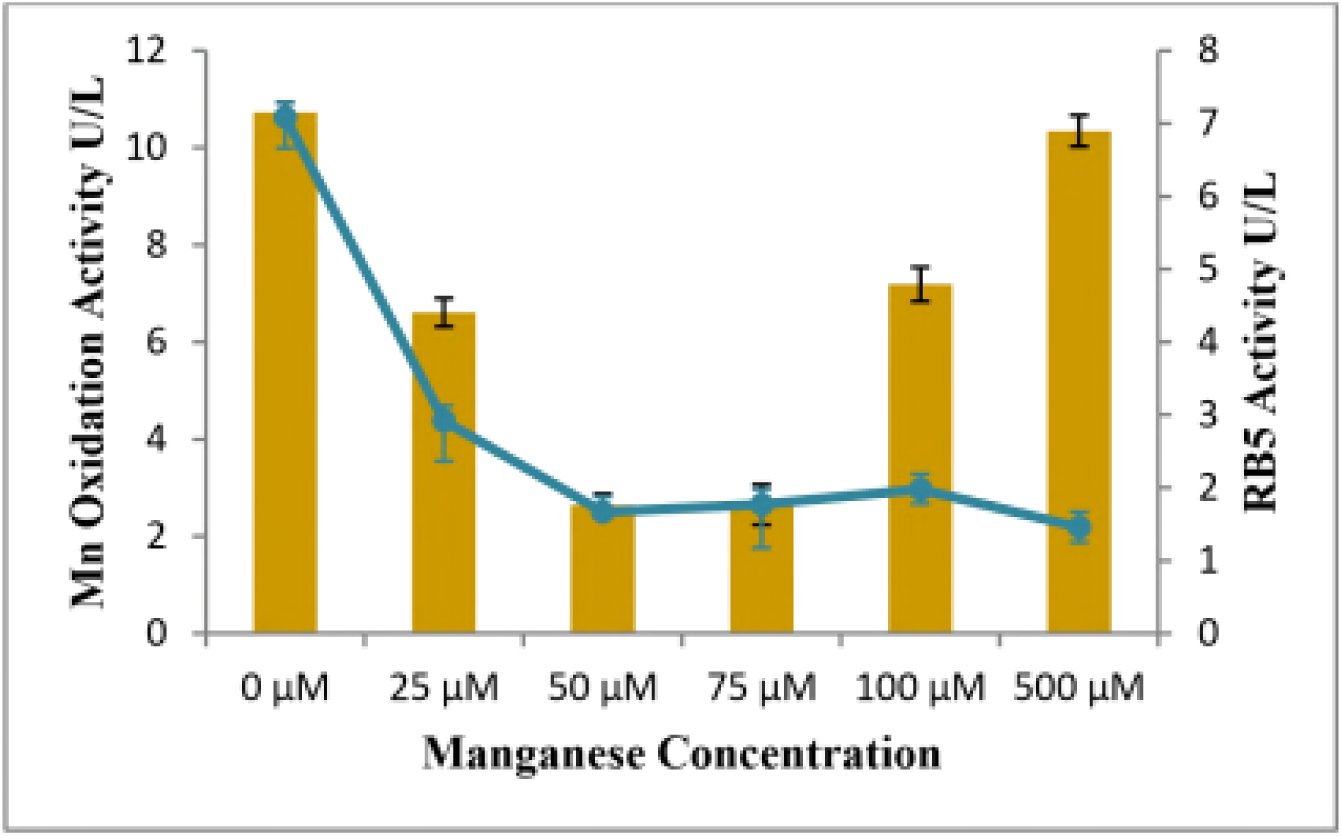
Manganese oxidizing peroxidase activity and RB5 decolorizing peroxidase activity at varying concentrations of manganese (0-500 µM).

**Fig 7:**
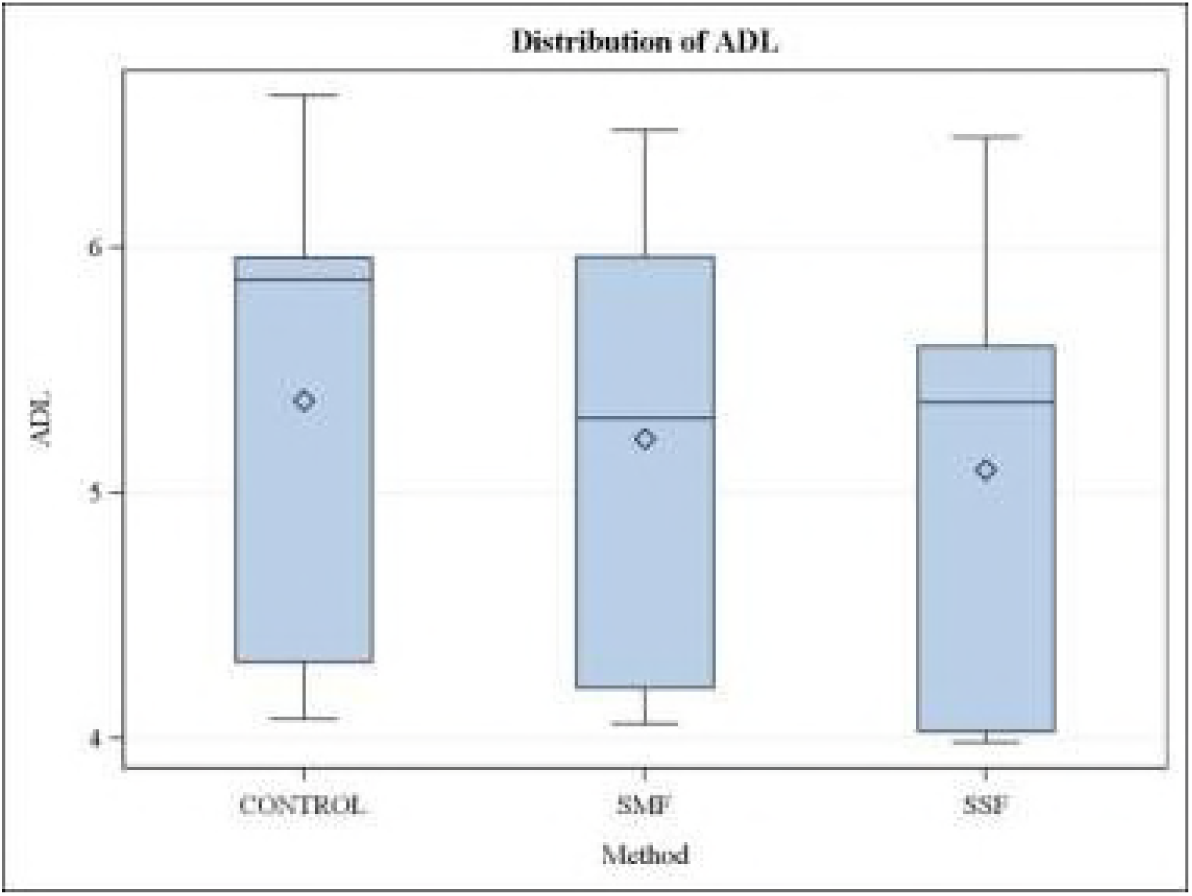
Distribution of ADL in control, solid state (SSF) and submerged (SmF) enzyme treatments of crop residues.

**Fig 8:**
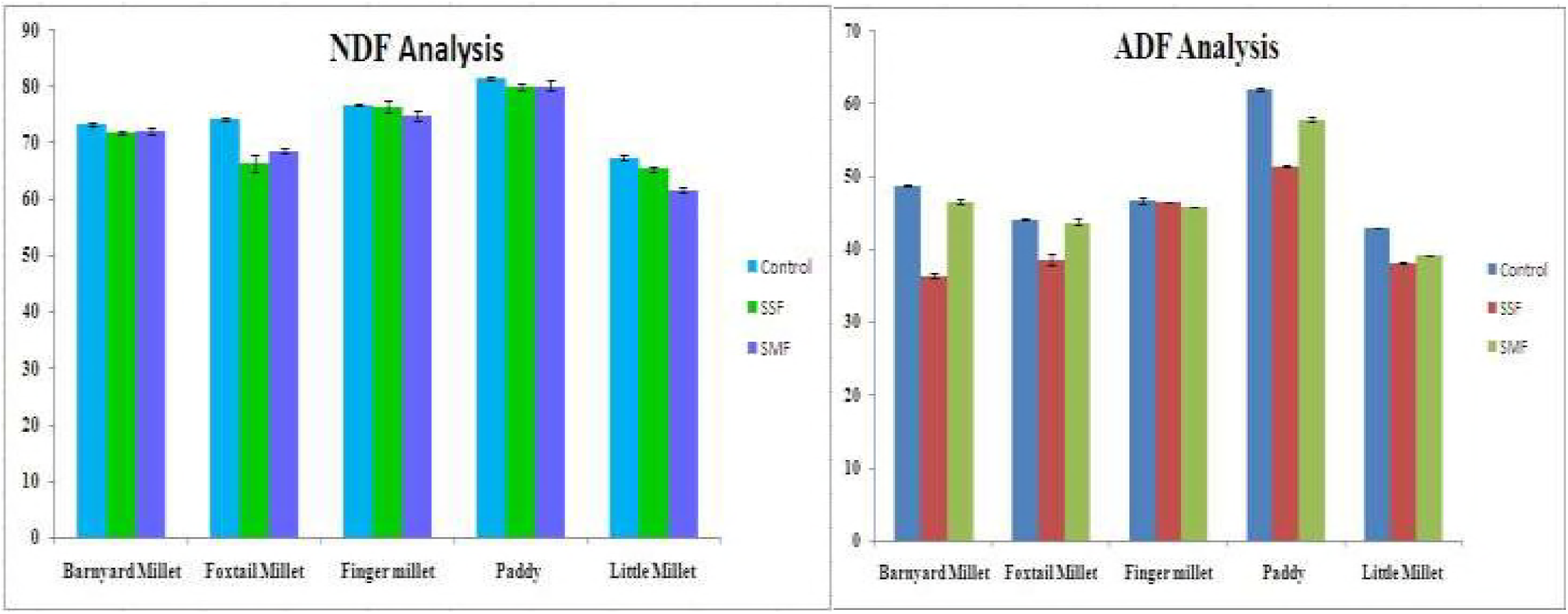
Distribution of NDF and ADF in control, solid state (SSF) and submerged (SmF) enzyme treatments of crop residues.

**Fig 9:**
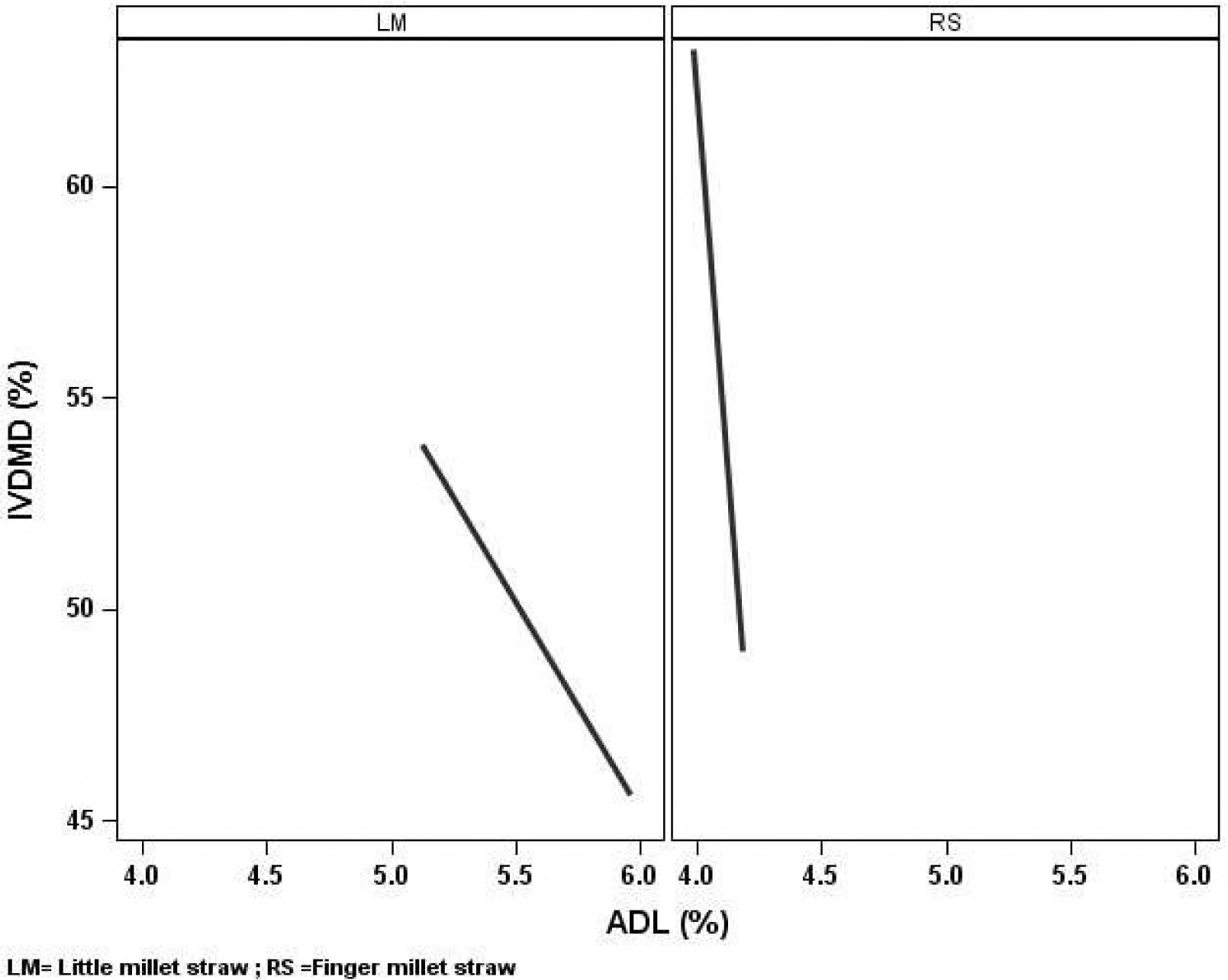
Correlation plot of ADL vs. IVDMD of solid state (SSF) and submerged (SmF) enzyme treatments of Little millet (LM) and Finger millet (RS) straw.

### Estimation of in vitro Dry Matter Digestibility (IVDMD)

Accretion of digestibility was observed with both little millet and finger millet straws subjected in the study. A maximum of 32% increase in digestibility compared to control was recorded with little millet straw treated with enzyme harvested through solid state fermentation and 29% for finger millet straw treated with the same enzyme. Enzyme produced through submerged fermentation provided 14% and 16% increase in digestibility for the former straws respectively (Table 2). Negative correlation between lignin content and digestibility is evident from the correlation coefficients -0.8 for finger millet straw and -0.46 for little millet straw. Augmentation of digestibility through delignification in the enzyme treated straws is illustrated in Fig 10.

**Table 2:** *In vitro* Dry Matter Digestibility (IVDMD) of control and enzyme treated straws.

| Straws | Method | IVDMD |
| --- | --- | --- |
| Little Millet | Control | 43.02±2.2 |
|  | SSF | 56.71±0.28** |
|  | SMF | 49.02±1.03 |
| Finger Millet | Control | 51.06±2.5 |
|  | SSF | 66.05±0.68** |
|  | SMF | 59.4±1.94 |

Values are means±SD (\* $P \leq 0.05$ , \*\* $P \leq 0.01$ , \*\*\* $P \leq 0.001$ )

## Discussion

Tannic acid and gallic acid are traditional substrates of Bavendamn reaction while guaiacol and syringaldehyde are phenolic lignin substructures produced upon lignin oxidation. These indicator compounds provided direct evidence of ligninolytic activity nevertheless diverse aromatic indicator compounds were used for substantiation of the lignin degrading ability of the isolates. The mode of wood decay by White-rot fungi in nature engages congregational action of two or more of the ligninolytic enzymes. Secondary compounds of this lignin macromolecule are readily oxidized by this ligninolytic machinery of the White-rots demonstrating their ligninolytic potential. Though these substrates are readily oxidized by the ligninolytic enzyme array, the mode of action and their potential differ. This is the rationale behind selective oxidation of certain substrates by the ligninolytic system. For this reason, Reactive black 5 (RB5) is reportedly oxidized by Versatile Peroxidase and Dye Decolorizing Peroxidases (5, 16). Its decolorization apparent for high redox potential peroxidases production was displayed by few isolates subjected in this study. The best isolate with this high potential for oxidation of RB5 was studied to decolorize azo dye RB5 and anthraquinone dye RBBR autonomously devoid of manganese. This strain designated as TAMI004 in the current study was relatively rapid and competent in its action on phenolics and dyes. This prompted us to characterize this strain further for its ligninolytic potential. Decolorization of Reactive Black started from day 3 and reached its maxima after 5 days and complete decolorization was observed after 7 days. Similar pattern of decolorization was also perceived with Remazol brilliant blue R, both dyes efficiently in manganese independent reactions. The oxidation of diverse substrates investigated places this enzyme in line with *Pleurotus* and *Bjerkandera* peroxidases (17, 18 and 19). White rot fungi typically oxidize natural lignocellulosic substrates through the collaborative effect of two or more of these ligninolytic enzymes. Here, growth conditions strongly influence the production of these enzymes from diverse eco physiological groups. We therefore studied the enzyme activities of this isolate in solid state and submerged fermentation modes. Activities of ligninolytic enzymes are reportedly affluent in solid state fermentation (20, 21) and likewise in our study, the activity of manganese oxidizing peroxidase was exponentially higher in solid state fermentation of wood than in submerged fermentation. In submerged fermentation, this manganese oxidizing peroxidase activity was significant in the medium without manganese. Though manganese oxidizing activity was predominant in media containing 0, 100 and 500 µM manganese, RB5 oxidizing peroxidase activity declined from 0 µM through 500 µM. This substantiates the existence of two distinct peroxidases in media void of manganese and in media with more than 100 µM manganese (22). Maximum activity peak in solid state fermentation was observed on day 12 after which activity declined and maintained till day 20, whereas in submerged fermentation activity peaked on 7^th^ day and dropped thereon. Maximal activity in submerged fermentation was observed at the end of log phase when the residual sugar concentration was almost 30%. Reducing sugar consumption was rapid during the log phase and was almost negligible during stationary phase. Preliminary investigation of the ligninolytic enzymes secreted by this wild isolate established the presence of high redox potential Versatile Peroxidase. Microscopic observation of spores, hyphae and its septa confirmed this strain as affiliated to the basidiomycetes group however, DNA based molecular identification techniques are sensitive and definite. For this reason, Internal Transcribed Sequence (ITS) region was acquired from the genomic DNA isolated from the fungus through the use of ITS 1 and ITS 4 primers. This strain was identified as *Lentinus squarrosulus* through comparison with BLAST of the fungal barcode sequence in a separate study undertaken in our laboratory. To the best of our knowledge, this is the first report describing production of Versatile Peroxidase from *Lentinus squarrosulus*. *Lentinus squarrosulus* is an edible White-rot mushroom of family Polyporaceae that grows extensively on decaying wood. Research analyses have ascertained the antioxidant and nutritional properties of this White-rot while ligninolytic abilities of this fungus need more investigation. Ligninolytic potential of this fungus was realized through the crude enzyme treated straws wherein there was a noteworthy improvement in the fiber fraction reduction as illustrated through proximate principles. The reduction in the lignin component of the crude enzyme treated straws is likely to improve its digestibility and concomitantly animal productivity. This was confirmed through the digestibility studies carried out on little millet and finger millet straws treated with the Versatile Peroxidase rich enzyme extract. Significant increase in digestibility in the enzyme treated straws confirms the potential of this enzyme to enrich the vastly available crop residues for utilization by ruminants. These results are salient from the ruminant nutrition perspective as apparently small increase in digestibility will matter extensively in livestock productivity. Though further research needs to be directed towards enhancing the activity of this high potential peroxidase and obtaining a deeper insight on its role in enhancing ruminant digestibility, the work is an impetus for further advancement in this direction. Production of high potential Versatile Peroxidase by this strain indicated through the current study will be of immense use in biotechnological and industrial applications especially in ruminant nutrition.

## Acknowledgment

The financial assistance by Department of Science and Technology (DST), Ministry of Science and Technology, Govt. of India, under the WOSA scheme (SR/WOS-A/LS-32/2016) is gratefully acknowledged by the first author. The authors thank the Director, ICAR - National Institute of Animal Nutrition and Physiology, Bangalore (Karnataka) India, for providing the necessary facilities to carry out the research work.

